# Airway epithelia clear rhinovirus infection via two waves of cell extrusion

**DOI:** 10.1101/2025.11.18.689008

**Authors:** Faith Fore, Dustin Bagley, Rocio Teresa Martinez-Nunez, Julia Aniscenko, Sebastian Johnston, Jody Rosenblatt

## Abstract

Epithelial barriers represent the first line of defence against pathogens, yet their role in innate immunity is typically relegated to pathogen detection and immune cell recruitment. This perspective ignores a fundamental evolutionary principle: epithelia defended against pathogens long before complex immune systems evolved. Here, we demonstrate that human bronchial epithelial monolayers retain this ancestral capacity, autonomously clearing rhinovirus (RV) within 24 hours by selectively extruding infected cells—a process we term virus-induced cell extrusion (VICE). VICE occurs in two waves: a rapid response occurring independently of virus entry, followed by a replication-dependent wave. Barrier-defective epithelia that cannot extrude fail to clear RV. While extrusion maintains barrier integrity and eliminates local infection, it also expels virus-laden cells, promoting transmission. Thus, VICE enables leukocyte-independent epithelial defence while inadvertently promoting viral transmission, reflecting an evolutionary strategy that prioritizes barrier integrity over containment. These findings redefine epithelia as central players in viral pathogenesis and host protection.

## INTRODUCTION

Rhinoviruses (RV) are non-enveloped positive-sense single-stranded RNA (ssRNA) viruses ^1–3^ and the primary cause of the common cold. Of three RV classes, RV-A and RV-B subdivide into major and minor groups, with the major group binding the intercellular adhesion molecule 1 (ICAM-1) receptor, the minor group binding the low-density lipoprotein receptor (LDLR), and RV-C binding to cadherin-related family member 3 (CDHR3) receptor ^2,4^. While often mild, RV infections can lead to severe respiratory complications in young children and individuals with asthma, chronic obstructive pulmonary disease (COPD), or cystic fibrosis.

Although RV primarily infects airway epithelia, the role of these tissues in immunity has often been restricted to that of an intermediary in signalling activation of innate and adaptive immune responses that recruit blood-based immune cells ^5,6^. However, airway epithelia are far more than passive signalling platforms: they form a critical physical barrier that filters pathogens and toxins from the environment, serving as a fundamental component of innate immunity ^2^. Epithelial cells are continually exposed to pathogens, pollutants, and irritants, and have evolved sophisticated mechanisms to maintain barrier integrity. We previously discovered a highly conserved, primordial process that drives most epithelial cell death called cell extrusion ^7^. During normal homeostasis to maintain optimal cell density, physical crowding triggers live cells to extrude by activating Piezo1 to form and contract a basolateral actin-myosin ring that seamlessly squeezes excess cells out apically ^8–10^. A Piezo-1-independent extrusion pathway also removes apoptotic or damaged cells, thereby preventing barrier function defects ^11,12^. Given the fundamental role of epithelia in protecting the body from environmental threats, we investigated how epithelial sheets cope with RV infections.

## RESULTS

### Human rhinovirus infection leads to extrusion of infected epithelial cells

To investigate how intact epithelial monolayers respond to RV infection, we infected human bronchial epithelial cells (16HBE14o^-^) with RVA01B or RVA02 (minor group), RVB14 or RVA16 (major group) at a <u>m</u>ultiplicity <u>o</u>f infection (MOI) of 3 for 24 hours. RV-infected cells were identified by immunostaining for the capsid protein VP3 (major group) or <u>d</u>ouble-<u>s</u>tranded RNA (dsRNA) (minor group), actin with phalloidin, and nuclei with DAPI. Confocal imaging revealed that both major and minor RV strains triggered extrusion of infected cells, characterised by a constricted actin-myosin ring beneath VP3^+^ or dsRNA^+^ cells (**Fig. 1a**). Thus, epithelial monolayers extrude RV infected cells and independently of host cell receptor, since all strains elicited the same response. Because RVA16, a major group, could be tracked from the earliest stages of infection using VP3 immunostaining, we used this strain for all future experiments.

**Fig. 1:**
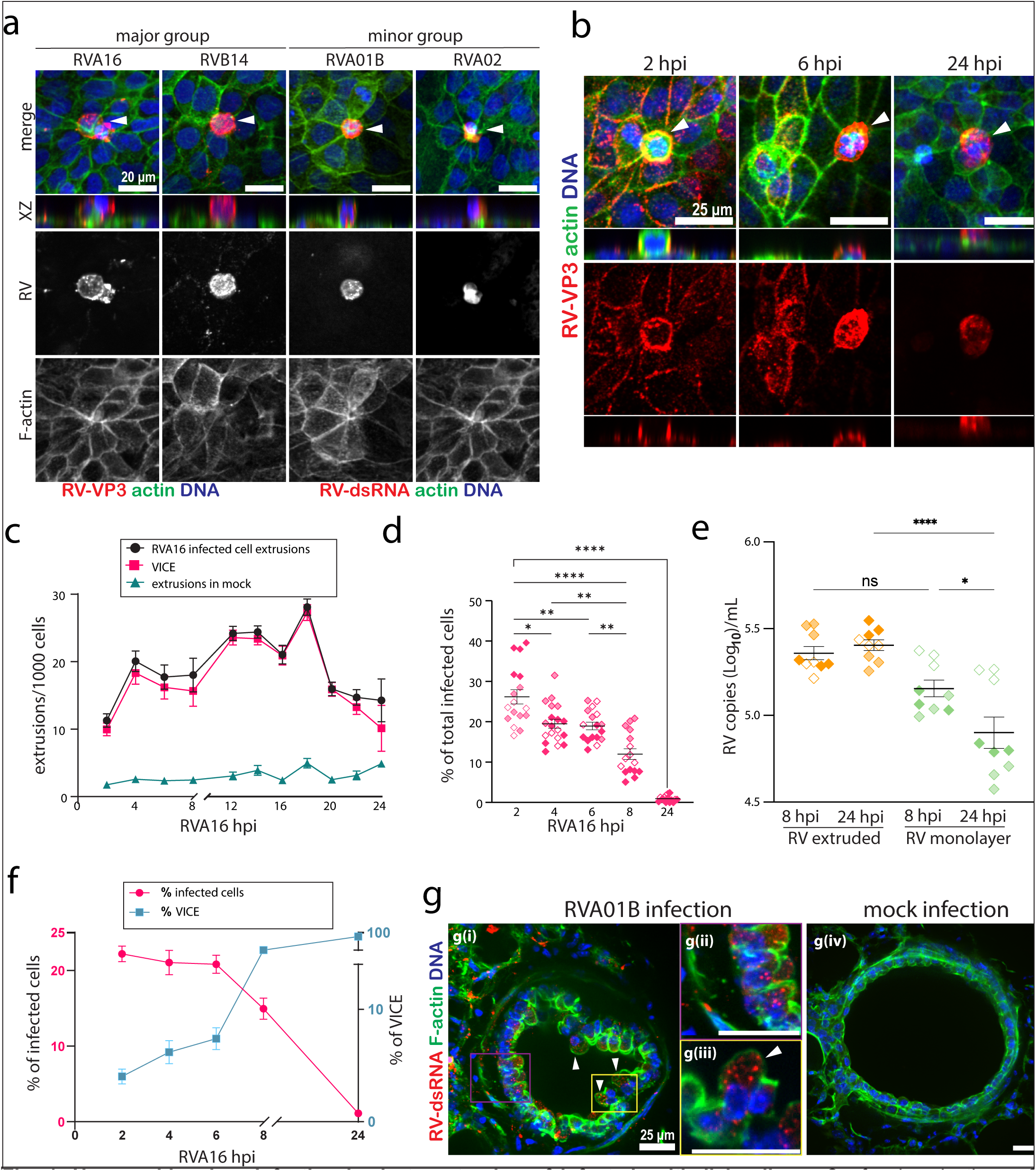
Human rhinovirus infection leads to extrusion of infected epithelial cells. **a**, Confocal projections of 16HBE14o^-^ monolayers infected for 24 hours with major-group (RVA16, RVB14) or minor-group (RVA01, RVA02) rhinoviruses and stained for VP3 or dsRNA, respectively, phalloidin and DAPI. **b**, Time course of RVA16 infection (MOI = 3) at 2, 6 and 24 h post-infection (hpi), quantified in **c**. **d**, Percentage of VP3+ cells remaining over time after RVA16 infection. **e**, RV RNA levels in extruded and monolayer-associated cells at 8 and 24 hpi, normalised to GAPDH and SDHA. **f**, Extrusion rates versus percentage of VP3+ cells. **g**, Confocal projections of mouse lung slices infected with RVA01 for 24 h versus mock controls, immunostained with dsRNA, phalloidin, and DAPI. n ≥ 3 biological replicates, indicated by differentially shaded points, with mean and s.e.m. **** p < 0.0001, *** p < 0.001, ** p < 0.01, * p < 0.05 from Welch’s ANOVA with Games–Howell post hoc test (**d**) and Mann–Whitney test (**e**). White arrowheads indicate extruding cells.

Remarkably, extrusion of RVA16-infected cells was detectable at the earliest timepoint of 2 hours post-infection (hpi) (**Fig. 1b**), when the VP3 immunostaining appeared predominantly at the cell periphery, suggesting that viral attachment or early post-attachment events may be sufficient to trigger extrusion. By contrast, at 24 hpi, VP3 localised throughout the cell during extrusion. While some non-infected cells extrude, as expected during homeostatic cell turnover, RVA16-infected VP3^+^ cells were preferentially extruded, since cell extrusion rates were significantly higher in infected monolayers compared to mock-infected controls (**Fig. 1c**). We termed this epithelial response to RV <u>V</u>irus-Induced <u>C</u>ell <u>E</u>xtrusion (VICE).

By 24 hpi, few VP3^+^ cells remained, steadily declining from ∼25% at 2 hpi to < 5% at 24 hpi (**Fig. 1d**). Additionally, quantification of RV RNA by RT-qPCR in both extruded and monolayer-associated cells (**Fig. 1e**) showed very few RV copies remaining in the infected monolayer by 24 hpi. Notably, both extruded and monolayer cells had equal RV RNA levels at 8hpi, indicating that roughly half of the virus had been cleared by VICE by this time, consistent with immunos-taining data in **Fig. 1b** and **1d**. While RT-qPCR measures total viral RNA and immunofluorescence enables quantification of the percentage of infected cells, both methods showed the same trend in viral reduction following RVA16 infection. Importantly, the percentage of infected cells progressively declined as the rate of extrusion increased (**Fig. 1f**), supporting a direct role of VICE in RV clearance. Thus, 16HBE14o^-^ cells eliminate RV infection through VICE in the absence of other innate immune cells.

To confirm if VICE occurs in real epithelial tissues, we infected ex vivo mouse lung slices with RVA01B, a minor RV strain that can infect mice. RVA01B induced high rates of extrusion in bronchiolar epithelia of mouse lung slices (at 24 hpi) (**Fig. 1g**). Because we lacked a structural RV antibody to RVA01B, we immunostained infected slices with dsRNA antibody at 24 hpi, at which point the extrusions were extensive enough to cause sheet detachment (**Fig. 1g(ii**)), similar to the damage resulting from broncho-constricted airways ^13^. The high extrusion rates may reflect a higher effective MOI, which is more difficult to calculate in tissue samples.

### An intact human airway epithelium clears rhinovirus-infected cells by VICE

As we found that extrusion alone could eliminate RV by extrusion, we investigated why this may not have been noted in previous studies. To do so, we infected BEAS-2B cells, a human epithelial cell line widely used in RV studies from normal human bronchial epithelial cells ^14^. Unlike 16HBE14o^-^ monolayers (**Fig. 1f**), BE-AS-2Bs could not extrude and failed to eliminate RV, resulting in RV accumulation over time (**Fig. 2a**). We reasoned that the inability of BEAS-2Bs to clear RV (**Fig. 2c**) might stem from their inability to form a tight epithelium with proper tight zonula occludins (ZO-1) and adherens junctions (E-cadherin) as seen in 16HBE14o^-^monolayers (**Fig. 2b**). Thus, although BEAS-2Bs derive from bronchial epithelia, they behave more like mesenchymal cells ^15^. To test if the inability of BEAS-2Bs to eliminate RV infection by extrusion was due to their lack of proper cell-cell junctions, we disrupted 16HBE14o^-^ junctions by calcium removal or chelation with EGTA (**Extended Data Fig.1**). Both ways of disrupting junctions prevented VICE-induced RV clearance, resulting in RV accumulation at rates seen in BEAS-2Bs at 24 hpi (**Fig. 2c&d**). These findings demonstrate that VICE-mediated clearance of RV requires an intact, extrusion-competent epithelial monolayer, revealing why studies using non-extruding cell lines may overlook this critical early antiviral defence **(Fig. 2e).**

**Fig. 2:**
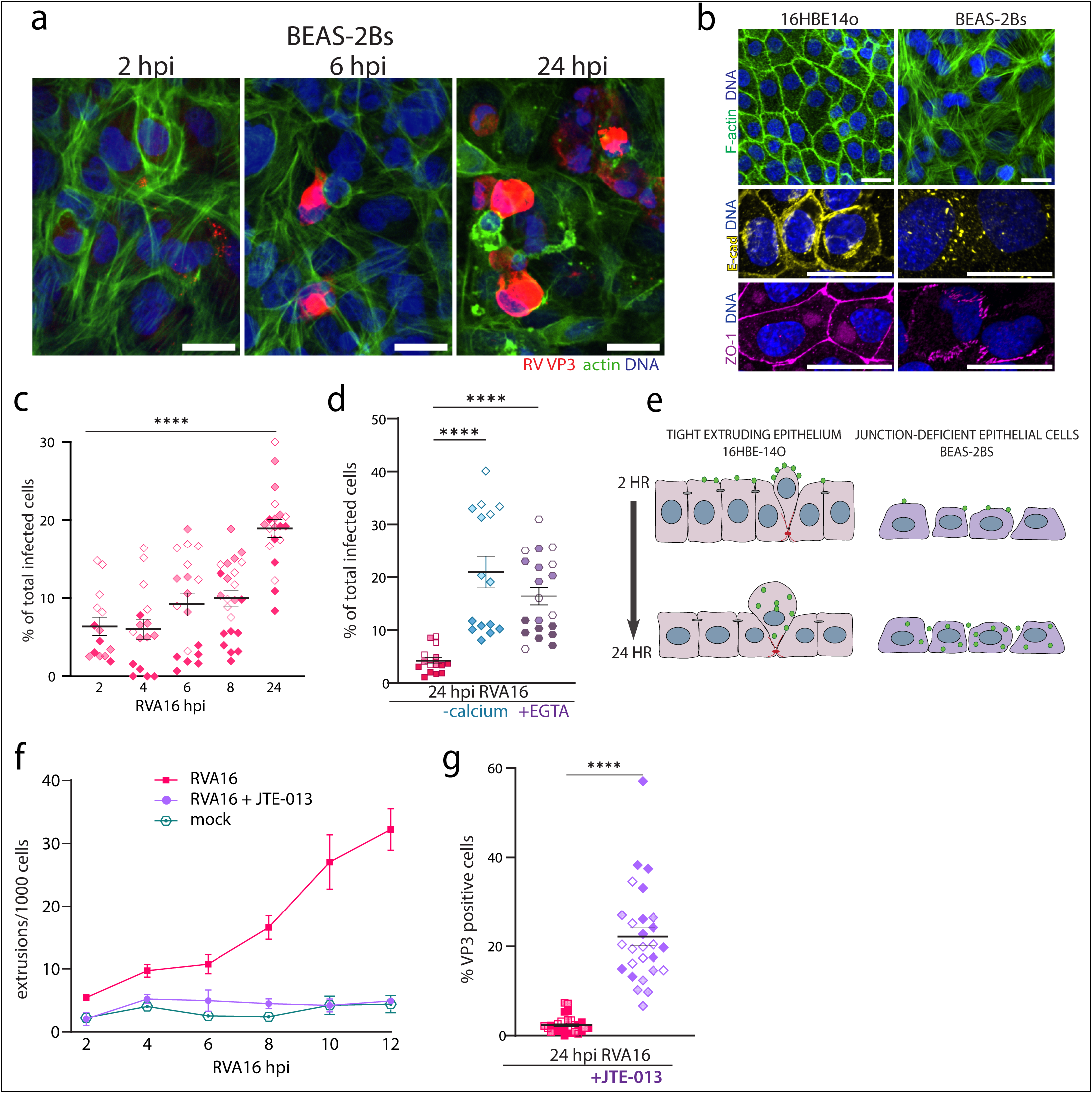
An intact human airway epithelium clears rhinovirus-infected cells by VICE. **a**, Confocal im-ages of Beas-2B cells infected with RVA16 for 2, 6 and 24 hours, scale bar=25um. **b**, Confocal projections of F-actin, E-cadherin and ZO-1 staining in 16HBE14o^-^ and Beas-2B monolayers, scale bar=10um. **c**, Percentage of VP3+ infected Beas-2B cells quantified every 2hrs after RVA16 in-fection. **d**, Percentage of VP3+ cells remaining at 24 hours in 16HBE14o^-^ monolayers with intact or disrupted junctions. **e**, Schematic illustrating clearance of infected cells in extrusion-competent (16HBE14o^-^) versus non-extruding (Beas-2B) epithelia. **f**, cell extrusion rates over time in RVA16-in-fected 16HBE14o^-^ monolayers ± JTE-013. **g**, Percentage of VP3+ cells remaining at 24 hours in RVA16-infected 16HBE14o^-^ monolayers ± JTE-013. n ≥ 3 biological replicates, indicated by differ-entially shaded points, with mean and s.e.m. **** p < 0.0001 from one-way ANOVA with Tukey post hoc test (**c**), Kruskal–Wallis test (**d and g**). RVA16 infections at an MOI of 3 unless otherwise indi-cated.

To confirm whether cell extrusion causes viral clearance, we blocked canonical extrusion signalling during infection. To extrude, a cell emits the lipid sphingosine-1 phosphate (S1P), which binds the G-protein-coupled receptor S1P receptor 2 (S1P_2_) to activate Rho-mediated contraction of an actomyosin ring basally and circumferentially that squeezes the cell out apically ^7,9,10^. Because our fixed, immunostained samples indicated that nearly all extrusions resulting from RV infection were RV-positive (**Fig. 1c**), we reasoned that we could score VICE rates over time using time-lapse phase imaging ± the S1P_2_ antagonist JTE-013. JTE-013 reduced VICE rates to the low basal rate seen in mock-infected cells (**Fig. 2f & Extended Data SMovie.1-3**), confirming that VICE requires the canonical extrusion signalling pathway. JTE-013 also blocked RV clearance from the monolayer at 24 hpi (**Fig. 2g**), confirming that RV elimination requires VICE. Together, these findings indicate that VICE eliminates RV-infected epithelial cells using the canonical extrusion pathway.

### RV induces two waves of cell extrusion, a live and an apoptotic one

While all apical extrusion pathways require the canonical S1P signalling tested above, different stimuli can trigger this signalling via different upstream pathways. To maintain homeostatic cell densities, most cells extrude while alive in response to crowding via Piezo1 ^8^, however, apoptotic stimuli can also induce extrusion to ensure that dying cells do not compromise the integrity of the monolayer ^7,11^. To investigate whether RV induces apoptotic or live-cell extrusion, we infected 16HBE14o^-^ monolayers with RVA16 and immunostained for the viral structural protein VP3, active caspase-3 (a marker of apoptosis), and phalloidin (actin) to visualise extrusion rings. We found that VICE was predominantly non-apoptotic during the first 8 hpi, shifting to mainly apoptotic extrusions for the next 12 hours (**Fig. 3a&b**). These two separate waves of live and apoptotic extrusion likely account for the two peaks of total VICE over 24 hours, seen in **Fig. 1c**.

**Fig. 3:**
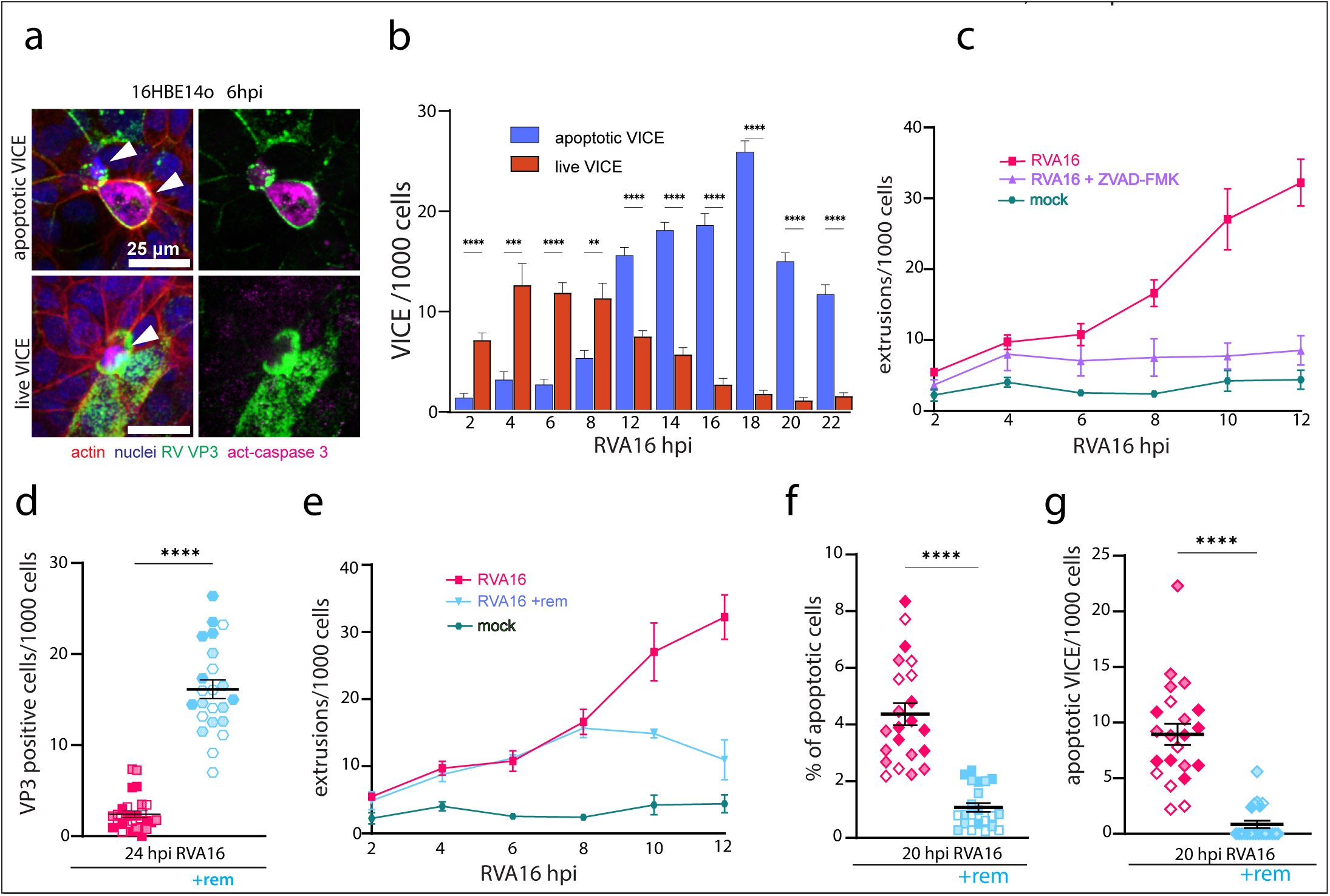
RV induces two waves of cell extrusion, a live and an apoptotic one. **a**, Confocal images of apoptotic and non-apoptotic extrusions at 6 h post infection, stained for actin (red), nuclei (blue), VP3 (green) and active caspase-3 (magenta). White arrowheads indicate extruding cells, quantified over time in **b**. **c**, Extrusion rates in control or z-VAD-FMK–treated RVA16-infected cells. **d**, Percentage of VP3+ cells at 24 h ± remdesivir. **e**, Extrusion rates ± remdesivir, every 2 h for 12 h by live-cell imaging. **f**, Apoptosis ± remdesivir. **g**, Apoptotic extrusions ± remdesivir. n ≥ 3 biological replicates, indicated by differentially shaded points, with mean and s.e.m. using an MOI of 3. **** p < 0.0001, *** p < 0.001, ** p < 0.01 by two-tailed t-test (**b**), Kruskal–Wallis test (**d**), Welch’s t-test (**f**) and Mann–Whitney test (**g**).

Given that apoptotic extrusion requires caspase activity ^11^, we tested whether the second wave of VICE re-quires caspases by treating infected 16HBE14o^-^ cells with z-VAD-FMK, an irreversible pan-caspase inhibitor ^16,17^. Time-lapse imaging revealed that extrusion rates were significantly reduced from 8 hours onward in z-VAD-FMK treated cells, compared to untreated controls, without impacting the first wave of extrusion **(Fig. 3c**). Since the apoptotic extrusion wave initiates at 8 hpi, the canonical time of RV replication^2^, we tested whether the second apoptotic wave of VICE requires RV replication using remdesivir, a cell-permeable mono-

### The first VICE wave is mechanical and independent of dynamin-dependent RV endocytosis

Since the first VICE wave is predominantly live cell extrusion, we tested if stretch-activated channels (SACs), which typically control live cell extrusion during homeostasis upstream of S1P2 signalling, control this first wave. To do so, we infected 16HBE14o^-^ in the presence or absence of gadolinium (Gd^+3^), a generic SAC blocker or GsMTx4, a tarantula venom peptide toxin that inhibits the transient receptor potential channels TRPC1, TRPC6, and Piezo1 ^20,21^. Both Gd^+3^ and GsMTx4 significantly reduced RV clearance by 24 hpi **(Fig. 4a)**, albeit to a lesser extent than blocking S1P with JTE-013. Live imaging demonstrated that both Gd^+3^ and GsMtx4 reduced VICE during the first 8 hours of infection (**Fig. 4b),** indicating that SACs control the first wave of VICE, accounting for reduced RV clearance. In both cases, GsMTx4 had a greater impact on VICE and virus clearance (**Fig. 4a&b**), confirming the link between VICE and RV clearance. These results confirm that VICE occurs in at least two waves, with the first live cell extrusion wave being controlled by SACs and the second apoptotic cell extrusion wave dependent on viral replication.

**Fig. 4:**
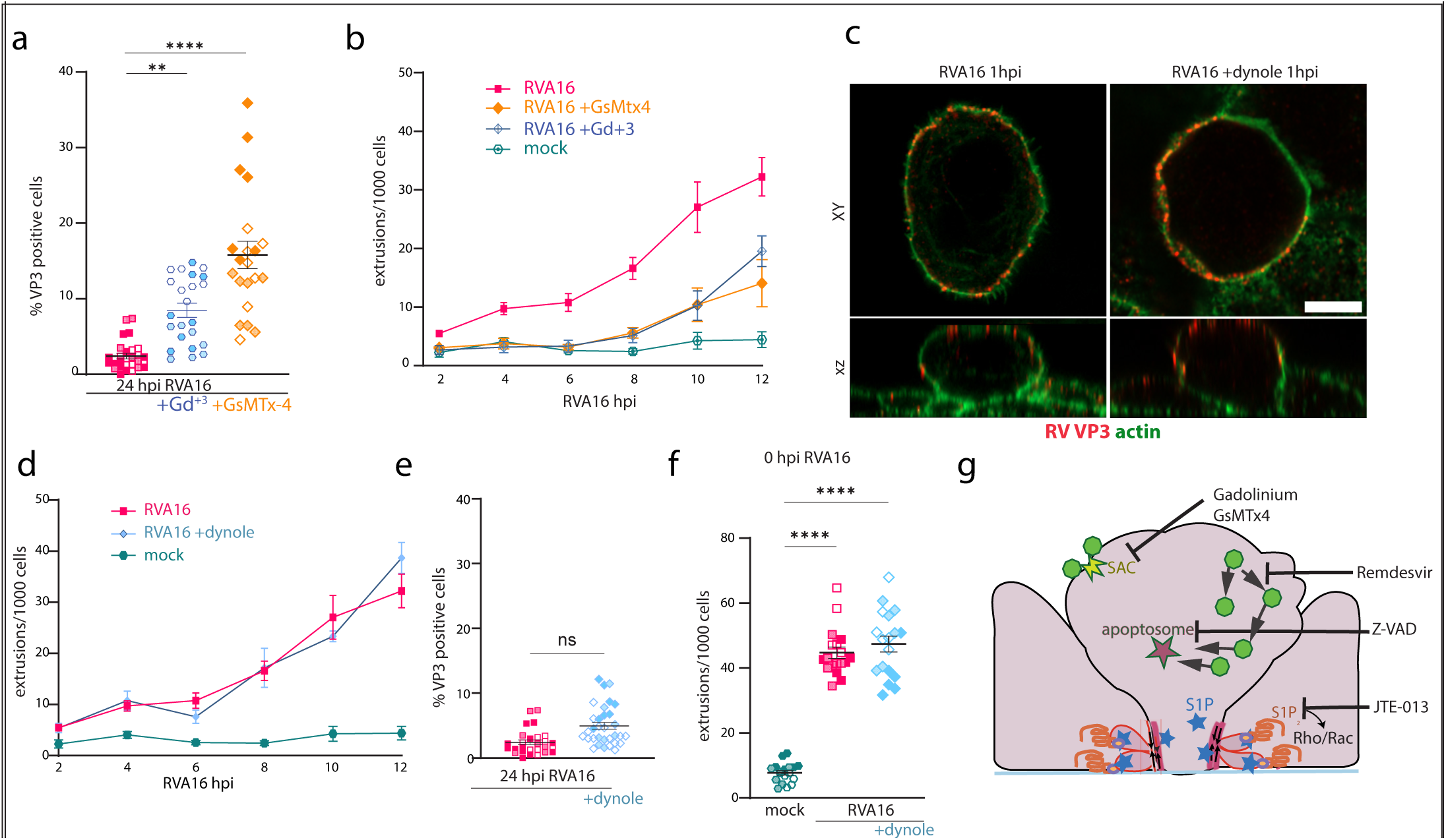
The first extrusion wave is mechanical and independent of dynamin-dependent RV en-docytosis or cell death. a, Percentage of VP3+ cells remaining at 24 hpi **±** Gd^+3^ or GsMtx4. **b,** Extrusion rates measured every 2 hours ± Gd^+3^/GsMtx4-treated monolayers. **c,** Confocal XY and XZ slices at 1 hpi **±** Dynole 34-2 (**bar=10 µm**). **d,** Extrusion rates ± Dynole 34-2 every 2 hours. **e,** Percentage of VP3+ cells remaining at 24 h **±** Dynole 34-2. **f,** Total extrusion events at 0 hpi **±** Dynole 34-2. **g**, Schematic showing inhibitors used in the study: Gd^+3^ and GsMtx4 block stretch-activated/TRP channels, z-VAD-FMK blocks apoptosis, remdesivir blocks viral replication and JTE-013 blocks S1P2 receptor. n ≥ 3 biological replicates, indicated by differ-entially shaded points, with mean and s.e.m. using an MOI of 3. ns p > 0.05; **** p < 0.0001; ** p < 0.01 by Kruskal–Wallis test.

Given that RV localised to the cell borders during VICE in the earliest extrusions measured (2 hpi, **Fig. 1b)**, we investigated if VICE might be activated before/during virus internalisation during the virus adsorption incubation step **(Extended Data Fig 3a)**. Using high-resolution confocal imaging, following a one-hour adsorption step at room temperature (see **Extended Data Fig 3a)**, we found all RV remained colocalised with cortical actin at the plasma membrane during VICE (Fig. 1c, n=8). To test if VICE requires RV up-take by receptor-mediated endocytosis ^22^, we infected 16HBE14o^-^ monolayers with RVA16 ± dynole 34-2, a cell-permeable allosteric inhibitor of dynamin-mediated endocytosis ^23^. A transferrin uptake assay confirmed that dynole 34-2 inhibits dynamin-dependent endocytosis in our system (**Extended Data Fig b)**. Indeed, we found the same peripheral VP3 staining in dynole-treated cells as seen in early VICE (**Fig. 4c**). The VICE scored during this early time frame where RV is restricted to the cell periphery could result from homeostatic background extrusions that had inadvertently been coated with RV. However, we found that the number of extrusions during RV absorption ± dynole was markedly higher than not only background mock controls but also than occurs during later VICE waves (**Fig. 4d**, ∼40 per 1000 cells versus ∼30 per 1000 cells at the peak, typically at 12 hpi). Thus, the early RV binding robustly promotes VICE. Additionally, inhibiting endocytosis with dynole 34-2 did not alter VICE rates over time (**Fig. 4e**) or significantly reduce RV clearance at 24 hpi, compared with control infections (**Fig. 4f**). While RV localises predominantly to plasma membrane during VICE between 2-8 hpi, the few extruding infected cells observed at 24 hpi had internalised virus (by VP3 staining) **(Extended Data Fig. 4d)**, potentially by macropinocytosis ^24^. Thus, epithelia use two strategies to ensure extrusion can eliminate RV infections: by activating live cell extrusion before the virus enters the cell, which is potentially mechanically activated, and by apoptotic cell extrusion in cells where RV has replicated (**Fig. 4g**).

### Extruded virus-infected cells can infect other cells

While epithelia eliminate RV by VICE, the cells that extrude are laden with virus, which could be highly infectious. To test if the cells extruded during VICE could act as ‘viral bombs’ that could infect other cells, we collected extruded cells at 16 hpi, washed them with medium, and added them to fresh BEAS-2Bs, HeLa-H1s, and 16HBE14o^-^ monolayers for 24 hours (**Fig 5a**). We found that extruded VICE cells could infect fresh monolayers by 6 hours post-VICE cell addition, and that this infection increased only in BEAS-2Bs and He-La-H1 cells, so that they were nearly all infected by 24 hours (**Fig. 5b&c).** By contrast, 16HBE14o^-^ eliminate RV by VICE by 24 hours, as expected (**Fig. 5b&c).** Notably, ∼100% of BEAS-2Bs were infected with viral bombs **(Fig. 5c)**, compared to only ∼20% with purified RV by 24 hours (**Fig. 2h**). While the infected extruded cells likely harbour significantly more viral units than cell-free virus, we cannot compare the actual MOI in each experiment, making it difficult to ascertain which are more infectious. However, our results reveal that viral bombs produced during VICE are competent to infect other cells and provide a more physiological route of virus spread than purified virus.

**Fig. 5:**
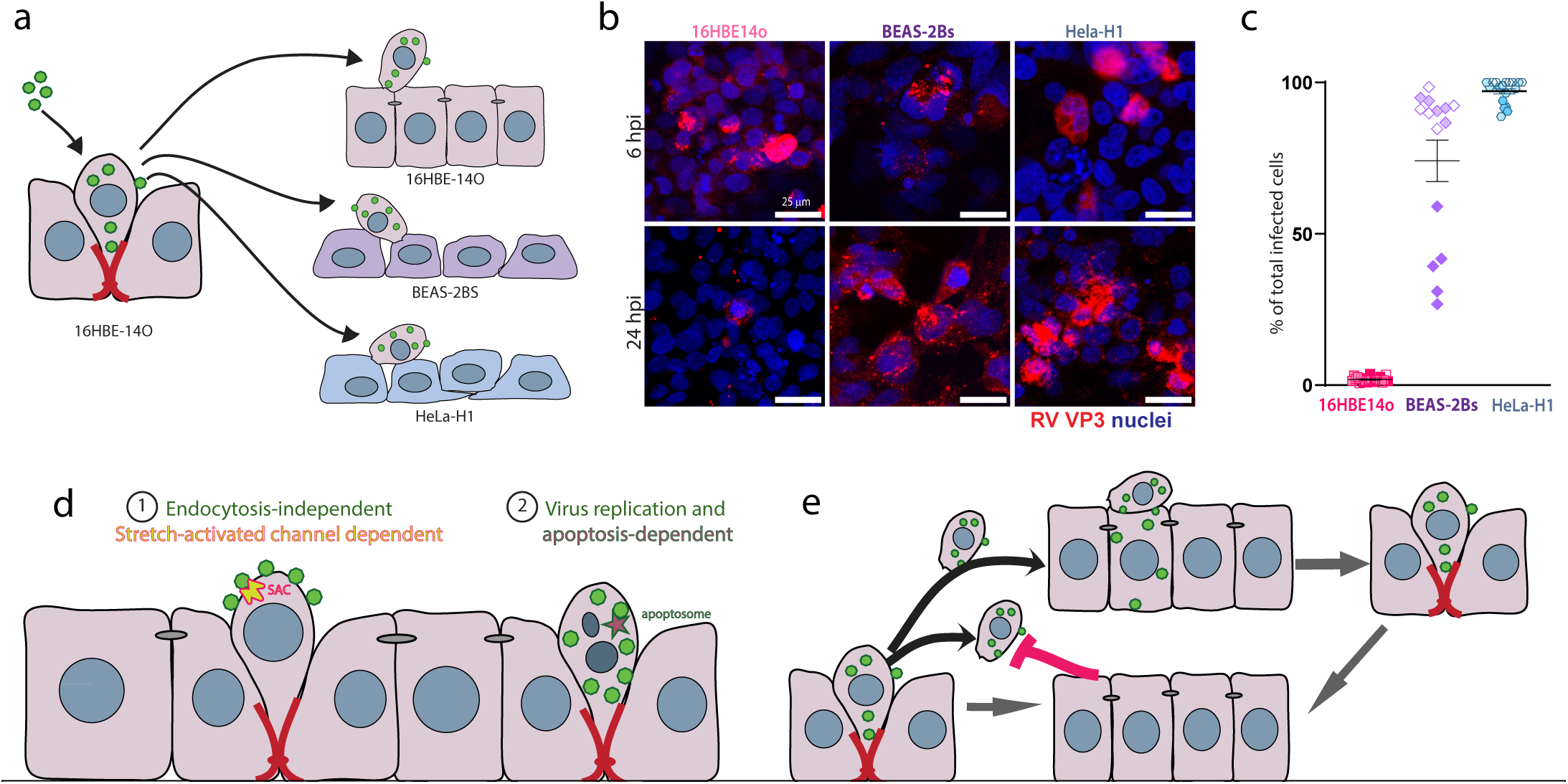
Extruded rhinovirus-infected cells can infect new monolayers. a, Schematic showing experimental design: extruded infected cells from RVA16-infected monolayers are added to naïve 16HBE14o^-^, Beas-2B, or HeLa-H1 monolayers. **b,** Confocal projections of each mono-layers at 6 h and 24 h after infected, extruded cell addition, quantified in **c. d,** Model illustrating that RVA16-infected monolayers clear infection via extrusion, and that extruded cells can ini-tiate infection in fresh naïve monolayers. n ≥ 3 biological replicates, indicated by differentially shaded points, with mean and s.e.m. using an MOI of 3. *** p < 0.001, **** p < 0.0001 by Kru-skal–Wallis test.

## DISCUSSION

Here, we found that epithelia rapidly clear rhinovirus by inducing virus-induced cell extrusion (VICE). VICE limits RV spread, providing a rapid protective response for airway epithelia without the need for other innate or adaptive immune cells. We believe VICE represents a primordial innate immune mechanism originating from many marine organisms, such as sea sponges, jellyfish, and corals, comprised of epithelia alone that have robust immune systems without the benefit of a blood system. We have identified two separate extrusion mechanisms that work in tandem to ensure the elimination of rhinovirus: an early endocytosis-independent wave of live extrusions, followed by a viral replication wave of apoptotic cells (**Fig. 5d**). Although highly efficient, the response is a double-edged sword, eliminating infected cells but facilitating rapid dissemination of infectious virus (**Fig. 5e**). Additionally, VICE in excess, as may be the case in asthmatics, could damage the epithelium, leading to an inflammatory response.

VICE-induced viral clearance is not limited to RV and likely represents the most common first step in innate immunity. Our findings support early observations using scanning electron microscopy showing that rhinovi-rus-infected bovine tracheal epithelia sloughed off, whereas those infected with parainfluenza did not ^25^. The increased severity of parainfluenza symptoms, compared to RV, may suggest that the inability to eliminate virus infections by VICE would lead to more systemic infections, needing other innate and adaptive immune responses. Conversely, the excess extrusion we noted in response to RV infections of mouse lung slices could trigger asthma attacks, as rhinovirus is the most common trigger of asthma exacerbations, whereas the lack of VICE response from parainfluenza may account for its link to milder asthma exacerbations ^26,27^. Similar extrusion or ‘sloughing’ has been noted in response to infection with RSV, enteroviruses, and *S. Tm* bacterial infections ^26–29^. Interestingly, RSV infection in infants, while common, can occasionally lead to excess sloughing of infected cells into airway lumens, causing obstruction and inflammation ^26^, and likely airway remodelling that can lead to asthma development ^13^. Thus, excessive or insufficient VICE can damage the airway barrier and inflammation, which could promote asthma, COPD, and cystic fibrosis down the line ^30–32^. Thus, VICE must be tightly regulated to strike a balance between viral clearance and tissue integrity.

Our findings on VICE have added to a growing body of research revealing different pathways that activate extrusion. Here, the first rapid pathway requires mechanosensitive channels, also implicated in enterovirus-induced extrusion ^29^. While we refer to this as a mechano-sensitive response, it may represent a non-specific irritant response akin to that from pollution, irritants, or allergens ^33,34^. Both GsMTX-4 or Gd^+3^, which block Piezo-1-dependent crowding-induced extrusion, can block transient receptor channels that trigger calcium influx in response to pollutants^21^ and might also trigger extrusion. Alternatively, virus binding Pattern Recognition or other receptors could induce mechanical membrane tension that activates the stretch-activated chan-nel-dependent wave, rather than crowding as seen during homeostatic live cell extrusion^8^. Viruses acting in a non-specific mechanical mode may explain why VICE occurs independent of host-viral receptor type (**Fig. 1**) and virus internalisation (**Fig. 4**), providing a generic response to new viruses or variants arising in the community. VICE clearance of RV before productive uncoating might contain viral spread or, on alternatively, create highly infectious vectors by promoting numerous viral particles to bind a single cell.

Further, our findings highlight the importance of viral replication in the second, apoptotic wave of extrusion that is triggered if virus-attached cells are not eliminated by the first rapid response. This could explain why remdesivir was not as effective as expected during the SARS CoV2 pandemic, if it also prevented virus elimination via VICE^35^. Additionally, because virus replication induces type I & III interferons (IFNs) that can trigger apoptosis, IFN activation may also control this second step^36^. Importantly, as asthma and COPD have deficient IFNs responses and apoptosis ^36–38^, VICE might be defective in asthma, leading to longer infections and more epithelial damage. Virus replication during the second wave of VICE may produce more infectious vectors upon cell rupture than those produced from the first wave.

Finally, our finding that extruded cells from VICE can efficiently infect other cells suggests two important points (Fig. 5e). One, extruded virus-laden cells can infect naïve monolayers efficiently, suggesting a physiologically relevant transmission route for viral dissemination. Two, while these viral bombs can infect new cells, the infected 16HBE14o^-^ monolayers do not appear to become reinfected from the viral bombs floating in their medium, suggesting a new form of epithelia-specific learned innate immune response, which will need further investigation. This finding makes practical sense as unchecked continuous reinfection and VICE within the same epithelia would rapidly destroy it. Identifying such a mechanism could lead to new therapeutic avenues aimed at preventing viral infections altogether, rather than merely managing their symptoms.

Clearly, understanding this ancient, intrinsic form of innate immunity will be of great importance to our management of virus spread and symptoms. We believe that using cell lines that do not model the tissues viruses normally encounter has led to oversight of this fundamental, primary step in innate immunity. Due to this oversight, the field has been overly focused on adaptive immunity, which has not been successful strategy for preventing RV spread. Understanding the mechanisms and signalling driving VICE has the potential to devise better prevention and management of RV and other viruses.

## ACKNOWLEDGEMENTS

We thank Michael J. Redd (University College London) for assistance with scanning confocal and helpful insight on the study, Pippa Hawes (The Francis Crick Institute) and Rui Galao Pedro for reviewing the manuscript, and members of the Rosenblatt laboratory for help with experimental design and figure layouts feedback.

## AUTHOR CONTRIBUTIONS

J.R., F.F. and D.C.B. conceived the study and designed the experiments. F.F. performed all experiments and data analyses. F.F. and J.R. interpreted the data and wrote the manuscript. All authors edited and approved the final manuscript.

## DATA AND MATERIALS AVAILABILITY

All data are available in the main text or the extended data. The source material link will be provided upon request.

## Extended Data Figure Legends

**Extended Data Fig. 1:**
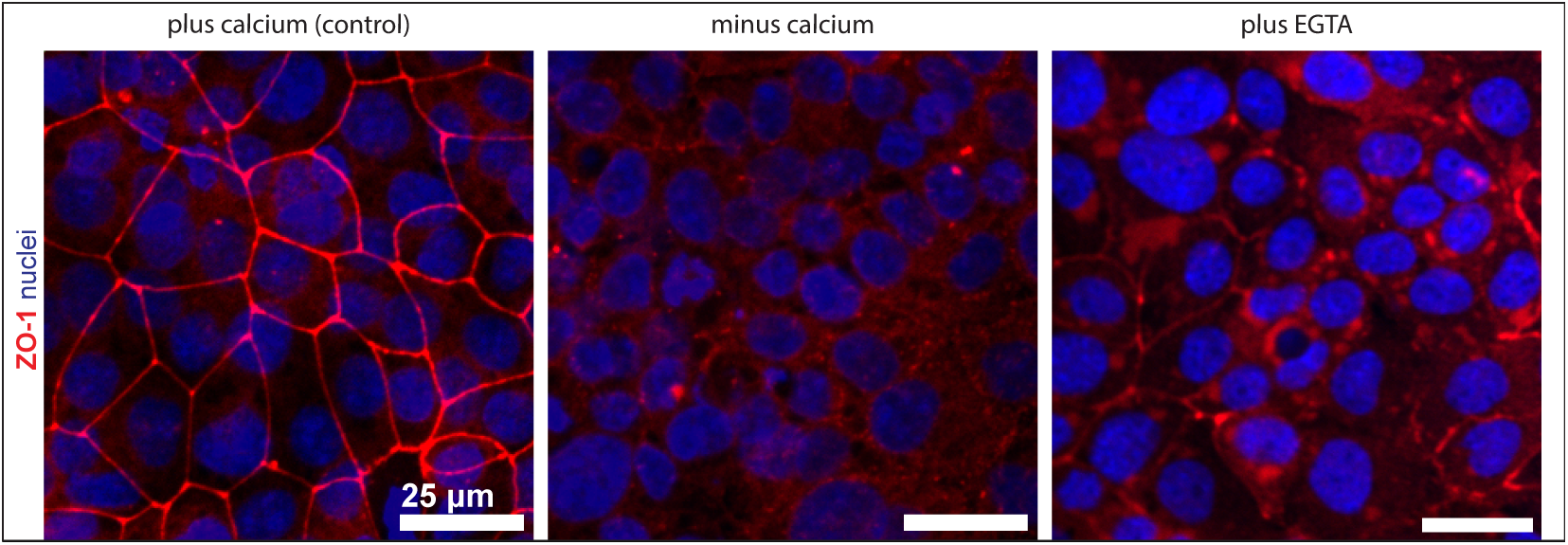
Disruption of tight junctions in 16HBE14o^-^ monolayers. Representative confocal images showing disruption of tight junctions in 16HBE14o^-^ monolayers by incubation in calcium-free medium or by chelating extracellular calcium with EGTA. Tight junctions were visualised using ZO-1 (zonula occludens-1) staining and nuclei with DAPI. Scale bar, 25 µm.

**Extended Data Fig. 2:**
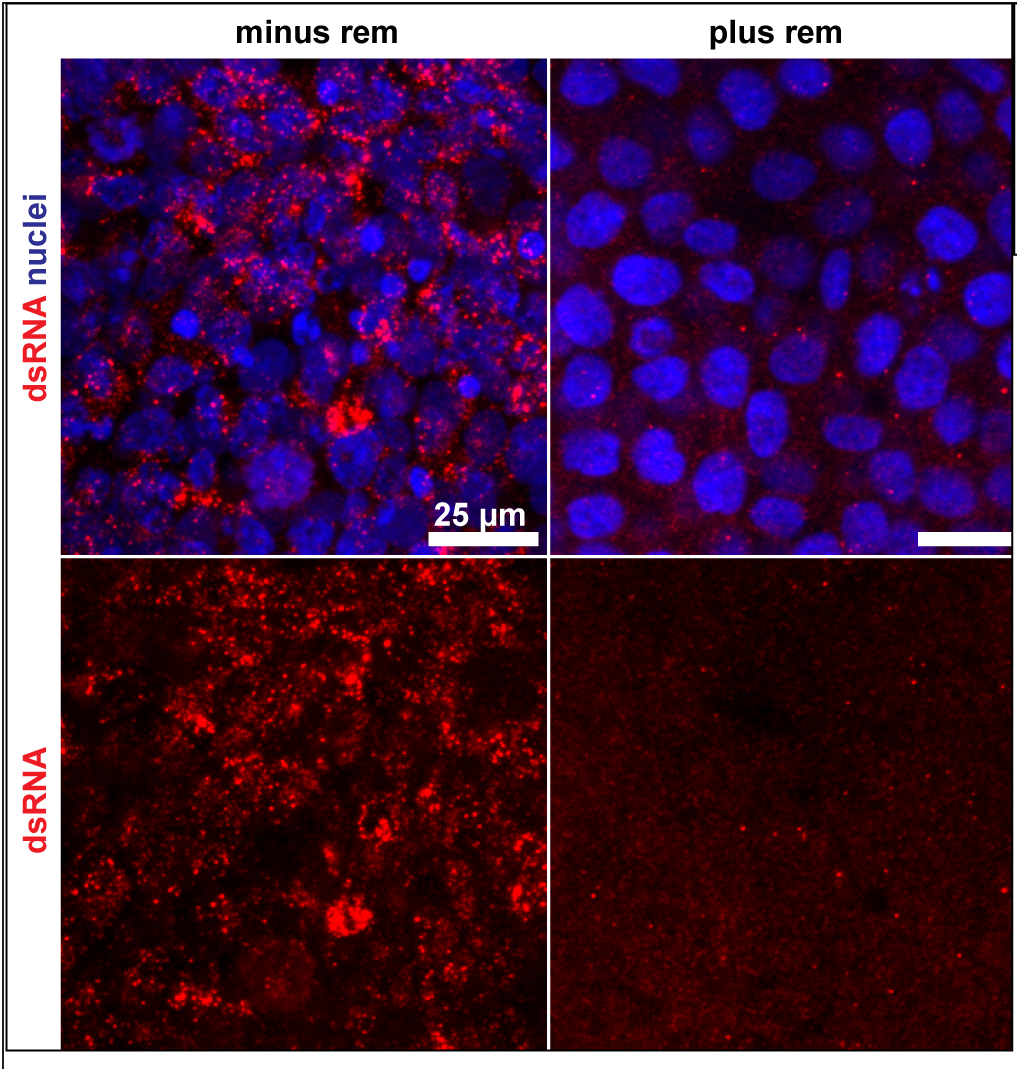
Remdesivir inhibits replication of RVA16 in HeLa-H1 cells. Representative confocal images of HeLa-H1 cells infected with RVA16 for 24 hours ± remdesivir Viral repli-cation was visualised by dsRNA staining and nuclei with DAPI (blue). Scale bar, 25 µm.

**Extended Data Fig. 3:**
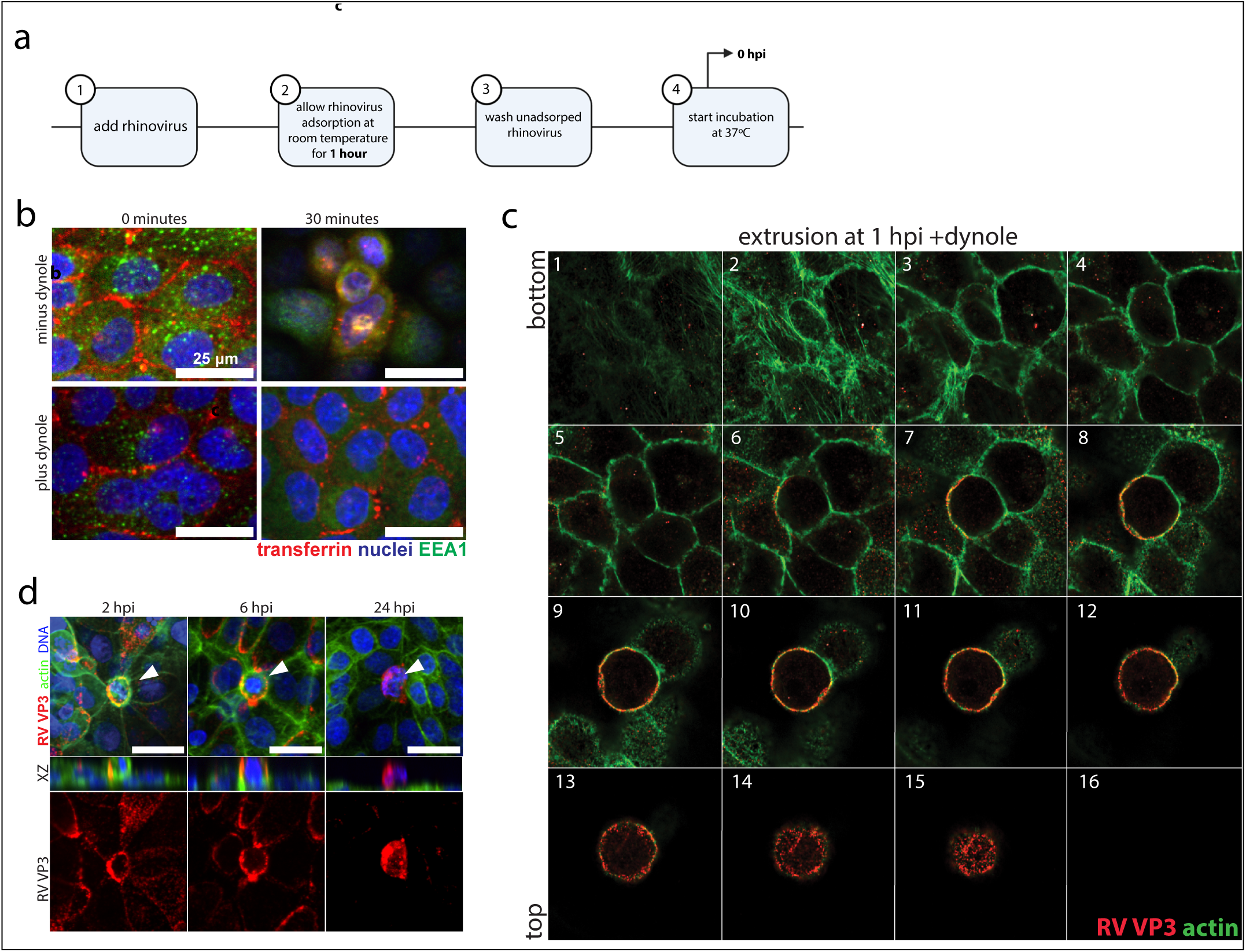
The early VICE wave is initiated during rhinovirus adsorption. **a,** Schematic of the infection experiment, indicating adsorption and infection steps (0 hpi). **b,** Representative confocal images showing transferrin uptake inhibited by Dynole 34-2, where early endosomes were also immunostained. **c,** Scanning confocal z-stack from basal to apical planes of a VICE cell at 0 hpi (one hour of RVA16 adsorption at room temperature). **d,** Representative confocal images of RVA16-infected cells ± Dynole 34-2. Scale bar, 25 µm.

**Extended Data SMovie1: Time-lapse imaging of mock-infected 16HBE14o^-^ monolayers.** Representative time-lapse sequence of mock-infected 16HBE14o^-^ monolayers imaged for 12 hours showing few extrusion events. Scale bar, 25 µm.

**Extended Data SMovie2: Time-lapse imaging of RVA16-infected 16HBE14o^-^ monolayers.** Representative time-lapse sequence of RVA16-infected 16HBE14o^-^ monolayers imaged for 12 hours, showing frequent extrusion events. Scale bar, 25 µm.

**Extended Data SMovie3: Time-lapse imaging of RVA16-infected 16HBE14o^-^ monolayers treated with JTE-013.** Representative time-lapse sequence of RVA16-infected 16HBE14o^-^ monolayers treated with the S1P2 receptor antagonist JTE-013, imaged for 12 hours showing extrusion events relative to untreated infected monolayers. Scale bar, 25 µm.

## METHODS

### Cell culture

16HBE14o (Merck, SCC150), HeLa-H1 (ATCC, CRL-1958) and BEAS-2Bs were grown in respective growth medium (**Table 1**) supplemented with 10% foetal bovine serum (FBS; 10270106) and 1% penicillin/streptomycin (P/S; 15070063) in a cell culture incubator at 37°C with 5% CO_2_. For all experiments, cells were seeded in growth medium and cultured until confluent. At confluency, the growth medium was removed, cells were washed twice with infection medium (**Table 1**) and incubated overnight in the same medium. All cell lines tested negative for mycoplasma contamination in periodic tests (Sartorius, 20-700-20).

### Rhinovirus stock production and titration

Viral stocks of RVA01B, RVA02, RVB14, and RVA16 (all from ATCC) were generated in HeLa-H1 cells and their titres were determined by measuring 50% tissue culture infectious dose (TCID_50_), as described previously ^51^. Briefly, cells at ∼90% confluency were washed twice with DMEM infection medium and inoculated with the virus diluted in infection medium. After 1 hour of gentle agitation at room temperature, cultures were incubated at 37 °C, 5% CO_2_ until ∼90% cytopathic effect (CPE) was observed. Cells were then dislodged, subjected to two freeze–thaw cycles at –80 °C, and clarified by centrifugation (4000 rpm, 15 min, 4 °C). The supernatant was filtered (0.2 µm; catalogue no. S2GPU01RE) and stored in aliquots at-80 °C.

For TCID_50_ determination, HeLa-H1 cells were seeded at 0.5 × 10^5^ cells/mL (150 µL/well, 96-well plate). Undiluted virus or 10-fold serial dilutions were added, and plates were incubated for 4-5 days at 37 °C, 5% CO_2_. CPE was scored microscopically, and TCID_50_/mL values were calculated using the Spearman–Kärber method ^52^.

### Rhinovirus infection of cell cultures

Cells were seeded at equal densities into 4- or 8-well IBIDI chambers or 24-well plates with size-1.5 coverslips and cultured at 37 °C with 5% CO_2_ until ∼95% confluency, followed by serum-starvation overnight in medium containing 2% FBS.

Drug cytotoxicity was evaluated using MTT (Merck, CT02) or CCK-8 (Merck, 96992) assays, according to the manufacturer’s instructions. For experiments with Remdesivir or Dynole 34-2, cells were pre-incubated with the drug for 2 h at 37 °C with 5% CO_2_ before infection. Viral stocks were diluted in infection medium to achieve a multiplicity of infection (MOI) of 3, with inoculum volumes adjusted for each format (150 µL for 24-well plates, 150 µL for 8-well chambers, 500 µL for 6-well plates, 200 µL for 4-well chambers). For drug treatments, the inoculum contained the drug at the corresponding concentration. Cells were inoculated for 1 h at room temperature with gentle agitation, washed three times with growth medium, and maintained at 37°C, 5% CO_2_ in fresh medium with or without drug.

Cultures were harvested at 0–96 hpi for downstream applications, including ELISA from supernatants, immunofluorescence (IF) from coverslips, or protein/RNA extraction. IBIDI chambers or glass-bottom 24-well plates were used for time-lapse imaging.

### Disruption of cell junctions Calcium free medium

Cells were seeded at equal densities and cultured in HBE growth medium without calcium (MEM, no calcium; Thermo Fisher Scientific, 11380037) supplemented with 2% Glutamax, 10% FBS, and 1% P/S for 36 h. The medium was then replaced with calcium-free HBE infection medium and incubated overnight. Infection was performed at MOI 3 in calcium-free medium for 1 h at room temperature with gentle agitation. Cells were washed three times and cultured in calcium-free growth medium at 37 °C, 5% CO_2_until harvest.

### Egtazic acid (EGTA) treatment

Cells were grown to ∼90% confluency and serum-starved overnight in infection medium. The following day, cells were pre-incubated in infection medium containing 1 mM EGTA. Infection was performed at MOI 3 in EGTA-supplemented infection medium for 1 hour at room temperature with gentle agitation. After inoculum removal, cells were washed three times and maintained in growth medium containing 1 mM EGTA at 37 °C with 5% CO_2_ until harvest.

Cells were grown to ∼90% confluency and serum-starved overnight in infection medium. The next day, cells were pre-incubated in infection medium containing 1 mM EGTA, then inoculated at an MOI of 3 in EGTA-supplemented medium for 1 h at room temperature with gentle agitation. Following inoculum removal, cells were washed three times and cultured in growth medium containing 1 mM EGTA at 37 °C, 5% CO_2_ until harvest.

### Immunofluorescence staining

Cells grown on coverslips were fixed with 4% paraformaldehyde (PFA) at designated time points and rinsed three times with DPBS. After blocking for 1 h at room temperature in IF blocking buffer (**Table**), cells were incubated overnight at 4 °C with primary antibodies (**Table 1**). Following three rinses and three 10-min washes in DPBS, cells were incubated with secondary antibodies (Table) for 2 h at room temperature. Nuclei were counterstained with DAPI (1:1000, v/v) for 10 min, washed, mounted in ProLong Gold (**Table 1**), and cured for 24 h before imaging.

For PCLSs, the procedure was identical, except secondary antibodies were incubated overnight at 4 °C, and washes were extended to three 30-min steps.

### Harvesting extruded cells

16HBE14o cells infected with RVA16 were processed for recovery of extruded cells. Culture medium was collected into a 15 mL Falcon tube, and the remaining monolayer was treated with 500 µL trypsin at room temperature for ∼2 min, with detachment monitored by light microscopy. Trypsinisation was quenched with 2 mL HBE infection medium, and detached cells were pooled with the collected medium. Suspensions were centrifuged at 1200 rpm for 5 min at 4 °C to pellet extruded cells, which were washed three times and resuspended in infection medium.

### Microscopy

Live-cell phase contrast and immunofluorescence imaging were performed on a Nikon Eclipse Ti2 microscope. For time-lapse phase imaging, a Plan Fluor 20× Ph1 DLL NA 0.50 objective, a Photometrics Iris 15 16-bit camera, and a CoolLED pE-4000 light source were used, controlled by NIS-Elements software (Nikon, v5.30.02). For immunofluorescence imaging, Plan Fluor 20×, 40× water-immersion, or 60× NA 1.40 oil-immersion objectives were used in combination with an iXon 888 Andor 16-bit camera, a Yokogawa CSU-W1 spinning disk confocal unit, and Toptica Photonics laser excitation, controlled by NIS-Elements (Nikon, v5.21.03).

### RNA extraction and cDNA synthesis

RNA from extruded or monolayer cells was isolated using TRIzol Reagent (Thermo Fisher Scientific) with minor modifications. Cells were washed with DPBS and lysed in 250 µL TRIzol per 1 × 10^5^-10^7^ cells. Chloroform (200 µL per 1 mL TRIzol) was added, and phase separation was performed by centrifugation (10,000 rpm, 30 min, 4 °C). The aqueous phase was transferred to tubes containing 1 µL glycogen blue (30 µg/mL), and RNA was precipitated with 0.5 vol isopropanol at-80 °C overnight. Pellets were recovered by centrifugation (10,000 rpm, 60 min, 4 °C), washed twice with 75% ethanol, air-dried, and resuspended in 20-50 µL nucle-ase-free water. RNA concentration and purity were assessed by NanoDrop spectrophotometry, and samples were stored at-80 °C.

cDNA was synthesised from total RNA using the reagents and volumes listed in **Table 1**. Reaction mixtures were incubated at 25 °C for 10 min, 42 °C for 60 min, and 70 °C for 10 min in a Bio-Rad T100 thermal cycler. cDNA was diluted with RNase-free water and stored at-20 °C.

### Quantitative PCR (qPCR)

qPCR was performed in 10 µL reactions containing 1 µL cDNA, using the ViiA7 RUO Thermal Cycler (Thermo Fisher Scientific) with specific probes and primers (**Table 1**). Cycling conditions were initial denaturation at 95°C for 1 min, followed by 40 cycles of 95 °C for 15 s and 60 °C for 30 s. Samples were run in duplicate, and amplification plots were used to confirm quantification within the exponential phase. Cycle threshold (Ct) values were averaged, with thresholds set within the linear lower third of each amplification curve. Target gene expression was normalised to the housekeeping genes GAPDH and SDHA, and presented as normalised RV copies.

### RVA01B infection of Precision Cut Lung Slices (PCLSs)

PCLSs were infected with RVA01B using a high virus titre of 5.5 × 10^7^ PFU/mL, corresponding to 1.65 × 10^7^ infectious units in the 300 µL inoculum applied per slice.

### Data quantification

Cellular events, including extrusions and total cell numbers, were quantified using NIS-Elements software (Nikon, v5.21.03). For time-lapse imaging, frames corresponding to 2 h intervals were analysed. Extruding cells were distinguished from dividing cells by the absence of daughter cell formation and by their floating morphology tethered to the monolayer. Cell counts were obtained using GA Analysis in NIS-Elements, with thresholds for brightness and size applied automatically and verified visually.

For fixed immunostained samples, quantification was performed by Z-stack inspection in NIS-Elements. Extruded cells were identified by actin rings at the basal plane and nuclei positioned above the monolayer. Nuclei were segmented automatically using GA Analysis, with segmentation accuracy confirmed visually.

Images were further processed using Fiji (ImageJ).

### Image and statistical analysis

Image presentation and statistical analyses were performed in GraphPad Prism (v10.3.1). Data represent the mean ± SEM from at least three independent biological replicates (n = 3). Statistical differences were evaluated using independent t-tests or one-way ANOVA with Dunnett’s or Tukey’s post-hoc tests, as specified in figure legends.

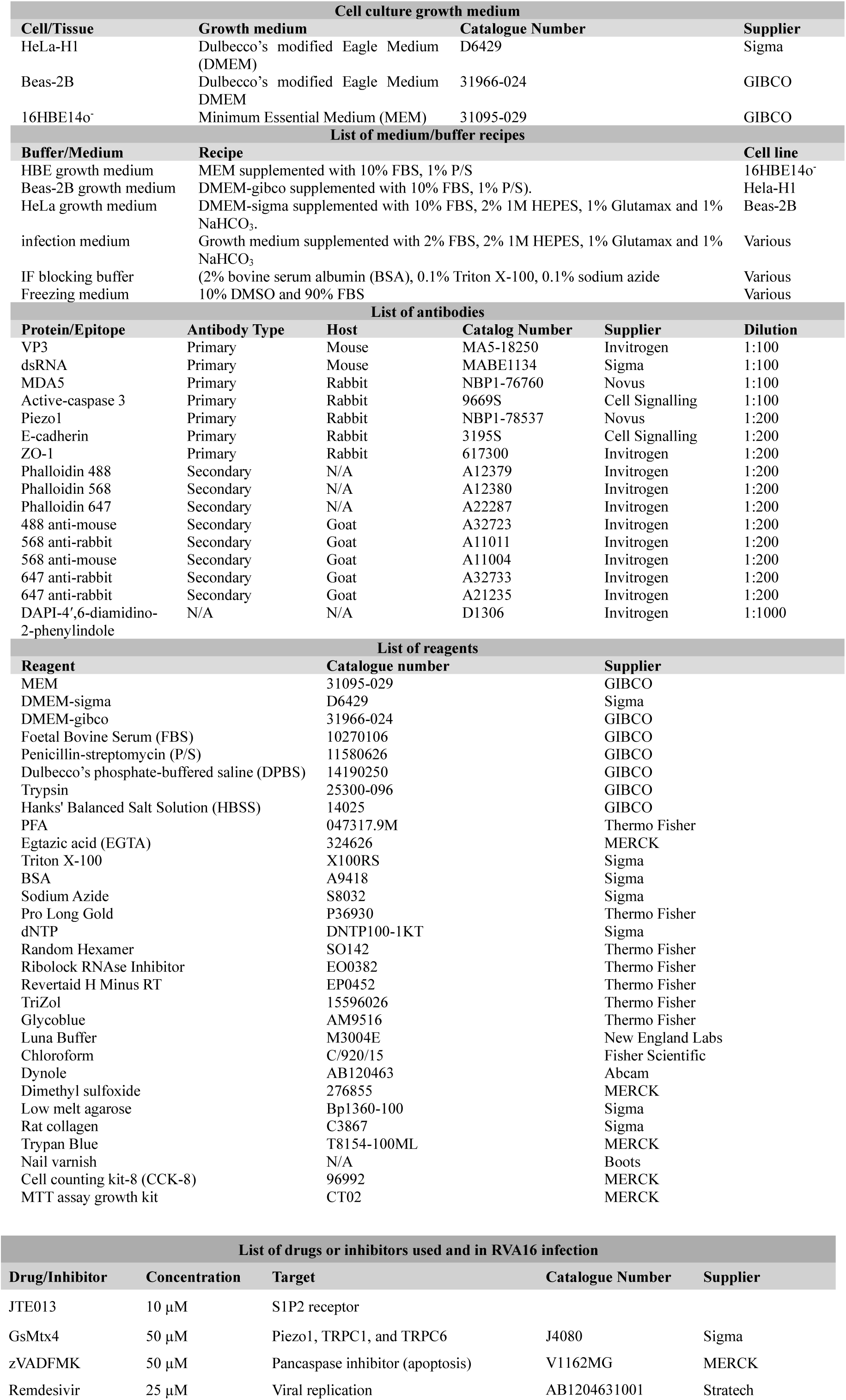
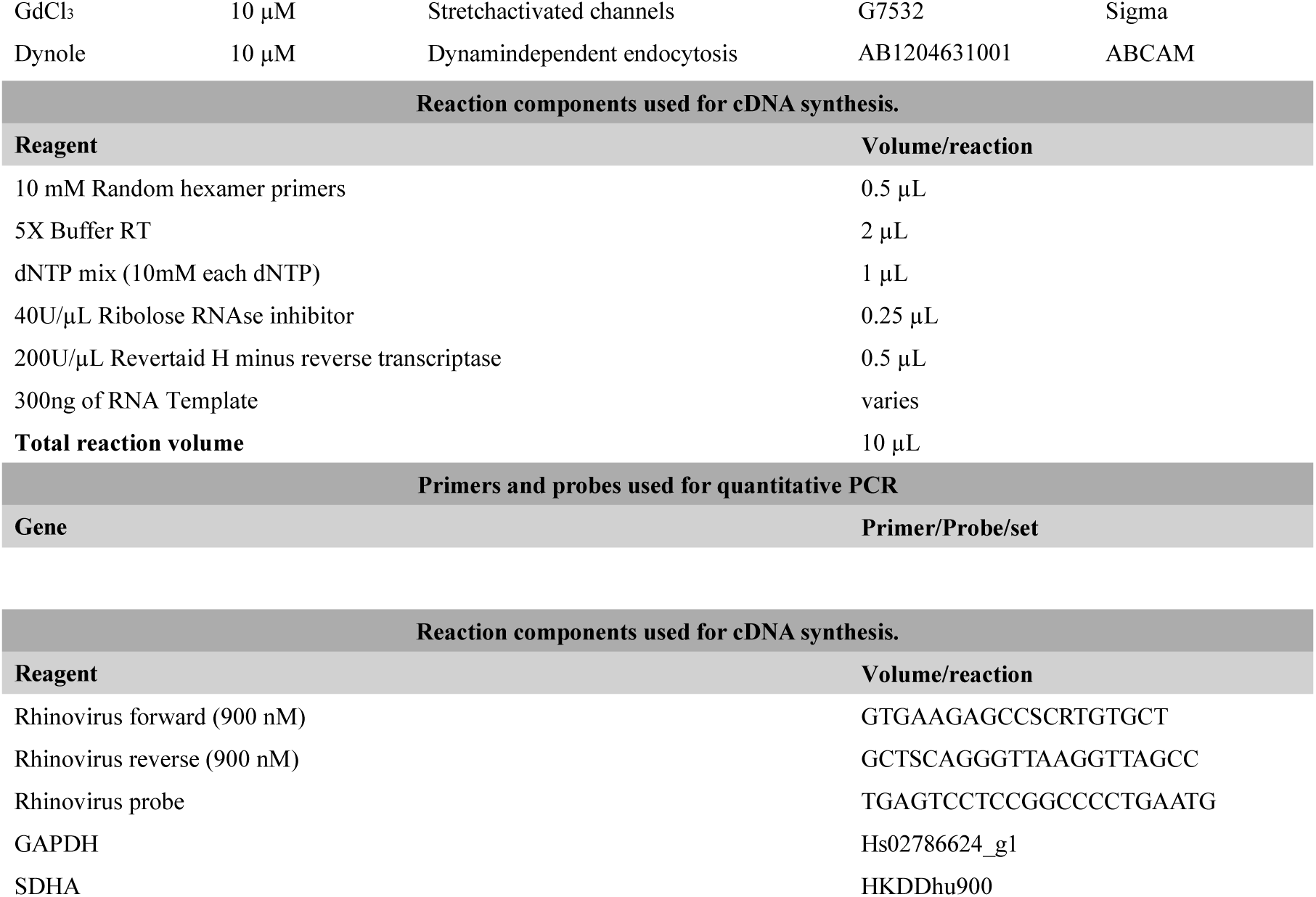

